# Proliferating cell nuclear antigen involves in temperature stress tolerance of *Ulva prolifera*

**DOI:** 10.1101/2023.02.17.529005

**Authors:** Hongyan He, Juanjuan Yang, Yuan He, Zhiyong Li, Caiwei Fu, Dongren Zhang, Mengru Li, Aiming Lu, Jingwei Dong, Jiasi Liu, Huiyue Gu, Songdong Shen

## Abstract

*Ulva prolifera* is the dominant species of “green tide”, and has higher tolerance to environmental stresses such as temperature. However, the molecular mechanisms are still unclear. Here, transcriptome analysis, Western blot and RT-qPCR analysis of *U. prolifera* suggested that, under temperature stresses (4°C, 36°C), the expression of *PCNA* and *CyclinA* was promoted, and the MAPK signaling was activated. Besides, the results showed that PCNA interacted with CyclinA. Interestingly, the expression of *miR-2916*, which was predicted to bind PCNA at -552∼-772, was negatively correlated with the expression of *PCNA* under temperature stresses (4°C, 36°C). In addition, the results showed that low temperature (4°C) had no obvious effect on the survival, the formation of cell walls, and the division of protoplasts. However, high temperature (36°C) had obvious effect on them. PCNA inhibitors increased the sensitivity of the protoplasts under temperature stresses. Together, our results suggested PCNA regulating the proliferation in response to the temperature stress of *U. prolifera* was associated with miR-2916/PCNA/CyclinA/MAPK pathway. In conclusion, the study preliminarily illuminates the molecular mechanism in response to temperature stress of *U. prolifera*, and may provide a new insight for prevention of green tide.

## Introduction

In the Yellow Sea of China, large-scale “green tide” causes series of deleterious impacts to the local ecosystem, since 2007(Anderson et al., 2012; Smetacek and Zingone, 2013; Hu et al., 2017; Blomme et al., 2021). *Ulva prolifera* (Ulvophyceae, Chlorophyta) has been proven to be the most dominant species during the form of “green tide” and self-reproduces in two forms: floating *U. prolifera* and attached *U. prolifera*(Cui et al., 2018; Wu et al., 2018a). Studies have reported that *U. prolifera* showed a high growth rate and huge biomass accumulation(Wu et al., 2018b; He et al., 2019). The growth rate of *U. prolifera* cultured at 25°C and 140 μmol/(m·s) reached 78.90% d^-1^(Bao et al., 2015; Hu et al., 2017.

*U*.*prolifera* growth is usually affected by various environmental stresses, such as temperature, light, and salinity(An et al., 2011; Fan et al., 2014). To adapt to these environmental stresses, *U. prolifera* has developed mechanisms to adapt to different types of stresses including high temperatures, cold, and high salt (McGlathery, 2001; Taylor et al., 2001). After exploring the key environmental factors that influence the growth and the reproduction of *U. prolifera*, researchers proposed that temperature was the main reason for the explosive proliferation of *U. prolifera*(Wang et al., 2015). The optimal temperature for the growth of *U. prolifera* is 20-27°C(Fu et al., 2008; He et al., 2017). Previous studies have reported that temperature affects photosynthesis, respiration, water balance, membrane stability and other metabolic processes of *U. prolifera*(Sousa et al., 2007; Luo et al., 2012; Zhang et al., 2013), how the molecular mechanism in response to temperature stress of *U. prolifera* still remains scarce.

In molecular biology, the proliferating cell nuclear antigen (PCNA) is considered to be a sliding clamp for DNA polymerase (Laquel et al., 1993). PCNA plays essential roles in eukaryotic cells during DNA replication, cell differentiation, senescence, and apoptosis(Prosperi, 2006; Bergink and Jentsch, 2009; De Biasio and Blanco, 2013). Previous studies have reported that PCNA binds with the S-phase-specific, CDK2-CyclinA complex(De Biasio and Blanco, 2013; Qian et al., 2019). Thus, it promotes the CDK2 binding to its substrate, thereby inducing the phosphorylation of the complex to regulate the cell cycle. PCNA also forms a stable quaternary complex with P21, CDK, and cyclins to regulate the apoptosis of cells(Xiong et al., 1992; Roig and Vázquez-Ramos, 2003; Raynaud et al., 2006). Gomez Roig et al. found that in maize plants, PCNA was bound to CDK cell cycle regulators, which were related to protein complexes. Moreover, it phosphorylated the DNA poIA gene (María de la Paz Sánchez et al., 2002). Thus, PCNA mediated the proliferation of maize crops. Raynaud et al. reported that Arabidopsis Tri-Thorax related protein 5 (ATRX5) is bound to PCNA through different subcellular localization mechanisms (Raynaud et al., 2006). Thus, it regulates the progression of cell cycle in Arabidopsis thaliana. In this study, we had to determine whether there is a PCNA protein in *U*.*prolifera*. Moreover, we also had to determine the role played by PCNA in response to temperature stress of *U. prolifera*.

Mitogen-activated protein kinase (MAPK kinase, MAPK) cascade plays an important role in environmental stress(Ichimura et al., 2000; Danquah et al., 2014; Bigeard et al., 2015). Environmental stress affects cell division and differentiation by activating the MAPK signaling pathway(Teige et al., 2004; Xing et al., 2008; de Zelicourt et al., 2016; Xu et al., 2018). However, the regulatory mechanisms of MAPKs in *U. prolifera* are not entirely known. Nearly 200 genes of small non-coding RNA (microRNA, miRNA) have been identified in animals and plants(Billoud et al., 2014; Tarver et al., 2015). In plants, some miRNAs were found to be involved in multiple stresses(Ju et al., 2017). For example, when the expression of miR319 was up-regulated, the resistance to cold stress was increased in *Arabidopsis*(Xin et al., 2010; Yang et al., 2013). However, the presence of miRNAs in algae was rarely reported.

In this study, we identified that PCNA regulated the proliferation of *U*.*prolifera* to acclimatize itself to temperature stress through miR-2916-PCNA-CyclinA-MAPK mediated pathway. The study used the molecular information in *U. prolifera*, and provided a better understanding of the thermotolerance mechanism in *U. prolifera* and a potential genetic target for prevention of green tide.

## Results

### Transcriptome analysis reveals that proliferation-related genes are involved in temperature stress resistance of *Ulva prolifera*

To understand the molecular mechanism in response to temperature stress of *U. prolifera*, we conducted transcriptome analysis of *U. prolifera* samples cultured at three different temperatures (UpTL, 4°C; UpTM, 20°C; UpTH, 36°C). In total, 129,396 genes were identified by implementing the Trinity method from the three libraries. All genes were performed differential expression analysis. The results shown that 6,304 differentially expressed genes (DEGs) were identified in UpTL vs UpTM. Among them, 3,180 genes were up-regulated, while 3,124 genes were down-regulated. There were 5,963 significant DEGs located in UpTH vs UpTM. Among them, 3,828 genes were up-regulated and 2,135 genes were down-regulated. There were 7,397 genes were up-regulated and 7,173 genes were down-regulated in UpTH vs UpTL (Figure 1A-C). In total, there was no significant difference in the number of DEGs under high and low temperature stress, suggesting that *U. prolifera* has a response to high and low temperature stress, but whether the mechanism is consistent or not still needs further analysis of the transcriptome. As shown in Figure 1D, there were 23,591 genes in UpTH vs UpTM and UpTL vs UpTM. The number of genes only responding to low temperature stress is 10,242, while the number of genes only responding to high temperature stress is 5113, further suggesting that the molecular mechanisms of *U. prolifera* responding to high and low temperature stress may be different.

**Figure 1.**
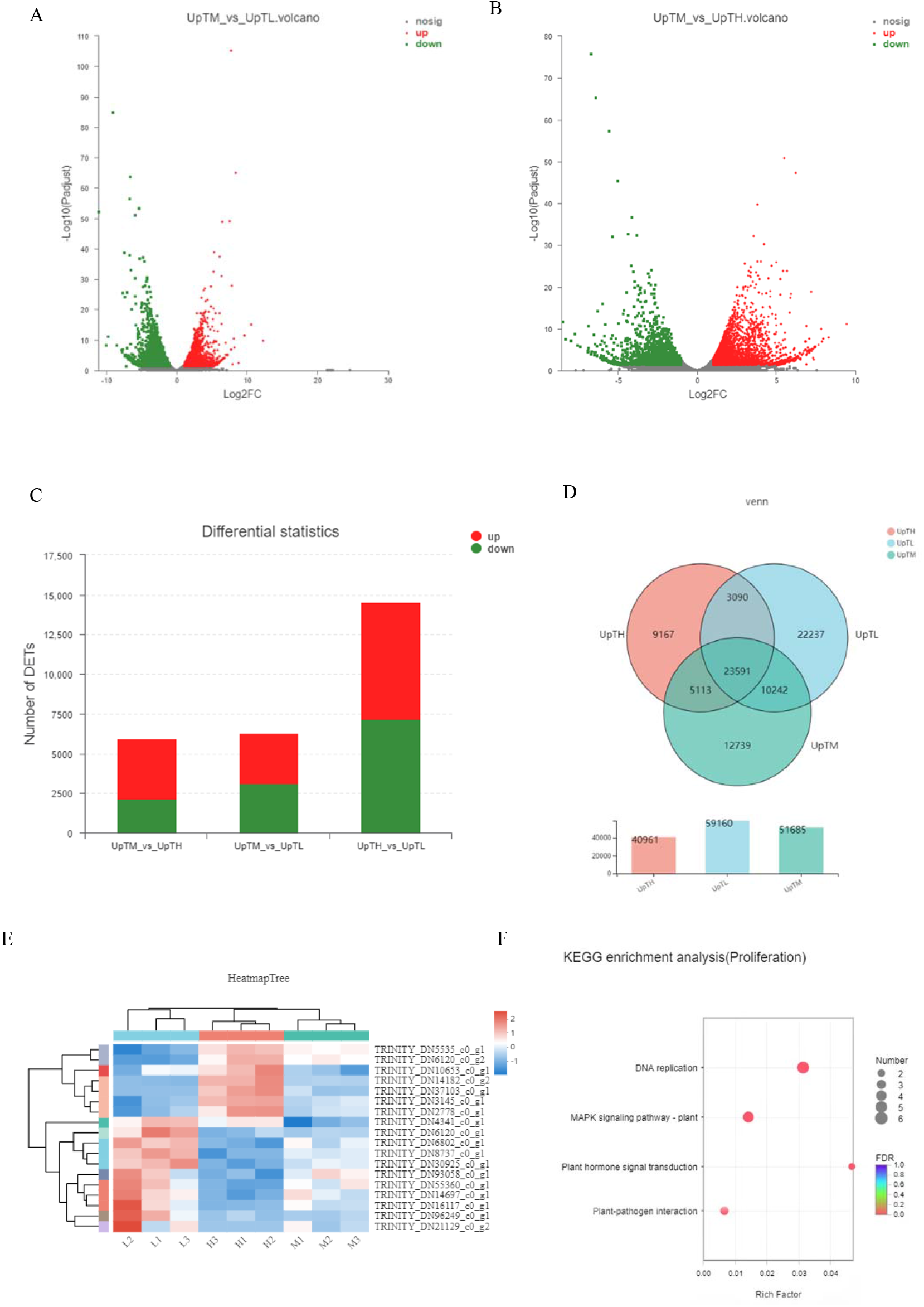
Transcriptome analysis of *Ulva prolifera* under different temperature. (A) Volcano plot of different gene between UpTL and UpTM groups. (B) Volcano plot of different gene between UpTH and UpTM groups. For volcanic plot, the horizontal and vertical axes represent the expression amount of genes; each point represents a specific gene or transcript; and the horizontal axis value corresponding to each independent point represents the expression amount of the gene in the control sample, and the vertical axis represents the expression amount of the same gene in the processed sample. In the figure, the red dots represent the genes that are significantly up-regulated, the green dots represent the genes whose expression is down-regulated, and the gray dots represent the genes whose expression is not significantly different among samples. (C) Statistics of differently expresses genes. (D) The Venn diagrams of differently expressed genes of *U. prolifera* under different temperature conditions. The circles with different color represent the total number of genes in *U. prolifera* under the same culture condition, and the intersection of the circles represents the number of genes expressed under each treatment condition. The abscissa represents the sample name, and the ordinate represents the number of genes. (E) Heat map of differential gene expression in *U. prolifera* under heat (H1, H2, H3) or cold (L1, L2, L3) stress compared with control (M1, M2, M3). The column represents the sample, and the row represents the gene. The color in the figure represents the levels of gene of each sample. Red represents the higher expression level of the gene in the sample, and blue represents the lower expression level. The left is the dendrogram of gene clustering, and the closer the two gene branches are, the closer their expression levels are; the dendrogram of sample clustering is on the top. The closer the two sample branches are, the closer the two branches are. The closer the expression patterns of all genes in a sample are, the closer the change trend of gene. (F) The annotation of KEGG pathway between UpTM and the temperature stress groups (UpTL, UpTH). The ordinate represents the KEGG pathway, and the abscissa represents the significance level of enrichment, which corresponds to the height of the column. The smaller the FDR, the greater the -log10 (FDR) value, the more significantly enriched in the KEGG pathway. UpTL (L), 4°C; UpTM (M), 20°C; UpTH (H), 36°C.

To explore the role of proliferation-related genes of *U. prolifera* in resisting temperature stress, we used the results of annotation of DEGs in NR database to search the annotated genes through the keywords: DNA replication and cell cycle. The results showed that there were 18 DEGs in UpTH vs UpTM and UpTL vs UpTM. The DEGs included cell division cycle (CDC2, CDC14, CDC7, PCNA), proliferating cell nuclear antigen (PCNA), cyclin, cyclin dependent kinases (CDKs), MCMs, and mitogen protein kinases (Figure 1E). Meanwhile, KEGG enrichment analysis was performed on the DEGs which associated with cell proliferation and division (Figure 1F). The results indicate that DEGs were mainly enriched in following processes: ONA replication, MAPK signaling pathway, plant hormone signal transduction, and plant-pathogen interaction.

### The expression analysis reveals that *PCNA* and *CyclinA* are involved in temperature stress resistance of *U. prolifera*

The DEGs related to cell proliferation in the sequencing results were compared with the genomic data of existing NCBI species. *PCNA* and *CyclinA* genes were found to be highly conserved with Arabidopsis, Rice, and other higher plants. The expression of *PCNA* and *CyclinA* in *U. prolifera* were studied to determine whether they were involved in cell proliferation in response to temperature stress. The samples were cultured at 4°C (UpTL), 20°C (UpTM), and 36°C (UpTH) respectively. The results showed that, the transcription level of *PCNA* and *CyclinA* were both up-regulated in UpTL samples than in UpTM sample. However, the transcription level of *PCNA* and *CyclinA* were up-regulated at UpTH on 2 days and 3 days, but were down-regulated after culturing the samples for 4 days (Figure 2A-F). The results suggested that low temperature stress induced the transcription of *PCNA* and *CyclinA*; short time high temperature stress induced them, but high temperature stress inhibited them over time. Western blot results showed that the protein level of PCNA increased, but the protein level of CyclinA increased after 3 days at UpTL. At UpTH, the protein level of PCNA and CyclinA increased on 2 days and 3 days, but the protein level of PCNA and CyclinA decreased after culturing the samples for 4 days. Likewise, the phosphorylation level of ERK (pERK) showed the same trend as PCNA (Figure 2G, H). This results indicated that low temperature stress up-regulated the protein level of PCNA and CyclinA; short time high temperature stress up-regulated them, but high temperature stress down-regulated them over time. And, the results also indicated that high temperature stress inhibited the MAPK signaling pathway over time; short-term high temperature stress and low temperature stress activated the MAPK signaling pathway.

**Figure 2.**
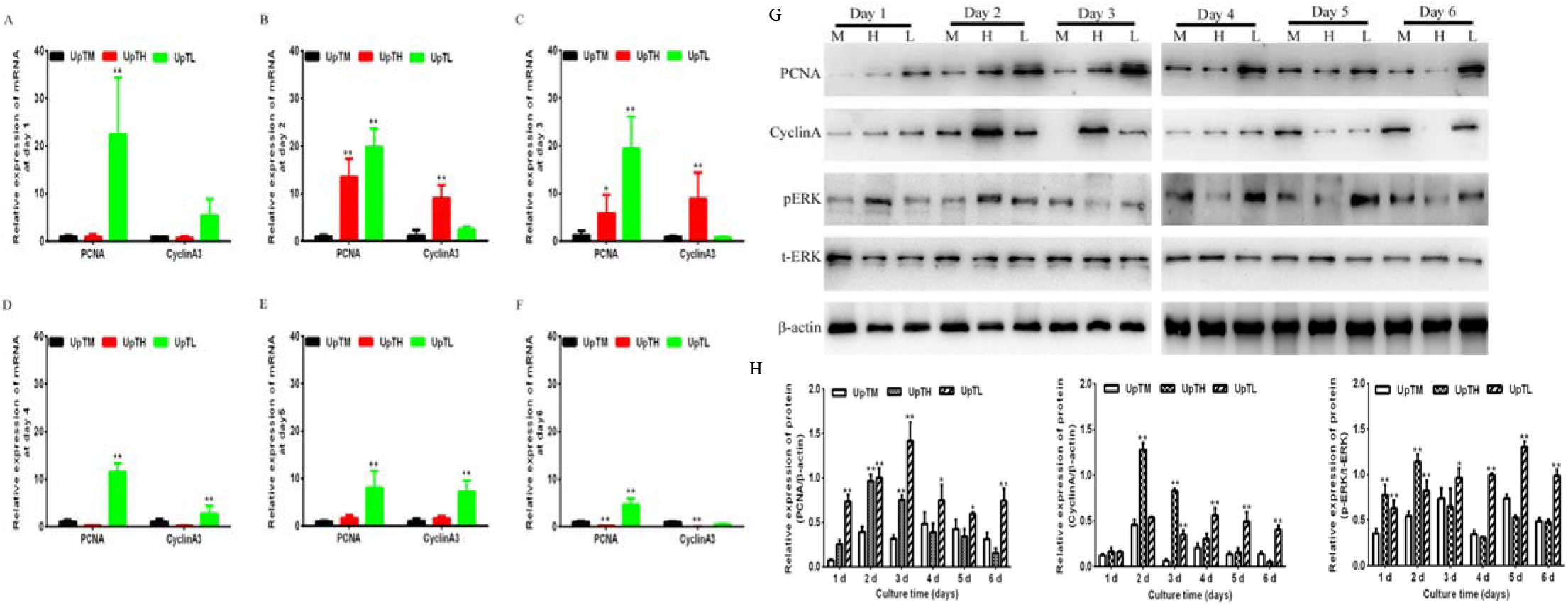
The levels of *PCNA* and *CyclinA* in *U. prolifera* cultured under different temperature at different time points. (A-F) The differential levels of *PCNA* and *CyclinA* in *U. prolifera* samples, which were cultured at different temperatures and different time-points. (G) Western blot analysis of PCNA, CyclinA, pERK, and tERK expression in *U. prolifera* samples, which were cultured at different temperatures and at different time points. (H) The density of the indicated molecular groups was analyzed by using ImageJ software. UpTM (M), 20°C; UpTL (L), 4°C; UpTH (H), 36°C; pERK, phosphorylated ERK; tERK, total ERK. *P<0.05,**P<0.01, vs. UpTM groups. The data were analyzed by performing one-way ANOVA (Tukey’s test).

### The expression analysis reveals that *miR-2916* is involved in temperature stress resistance of *U. prolifera*

Current studies have found that a variety of miRNA are involved in plant response to temperature stress by regulating the expression of target genes. In this study, based on the report (Huang et al., 2011), the conserved miRNA of *U. prolifera* was compared with that of other algae and plants using miRBase database. The results showed that *miR-2916* of *U. prolifera* had high similarity with that of *Populus euphratica*. At the same time, through psRNATarget and psRobot, we predicted the target gene of the miR-2916. The results indicated that a binding site is located between miR-2916 and -552∼-772 of PCNA (Figure 3A). Furthermore, we predicted that miR-2916 of *U. prolifera* and PCNA of *Chlamydomonas reinhardtii* had binding sites at positions -799∼-819 (Figure 3B). Thus, the combination of miR-2916 and PCNA is universal in the algae. Furthermore, we found that the expression of *miR-2916* decreased initially. Then, the transcription level of *miR-2916* up-regulated after 4 days in the UpTH; However, the transcription level of *miR-2916* was down-regulated at UpTL (Figure 3C-H). Therefore, the results showed that *miR-2916* involved in temperature stress resistance of *U*.*prolifera*, and the expression of *miR-2916* was negatively correlated with the expression of *PCNA* under temperature stress.

**Figure 3.**
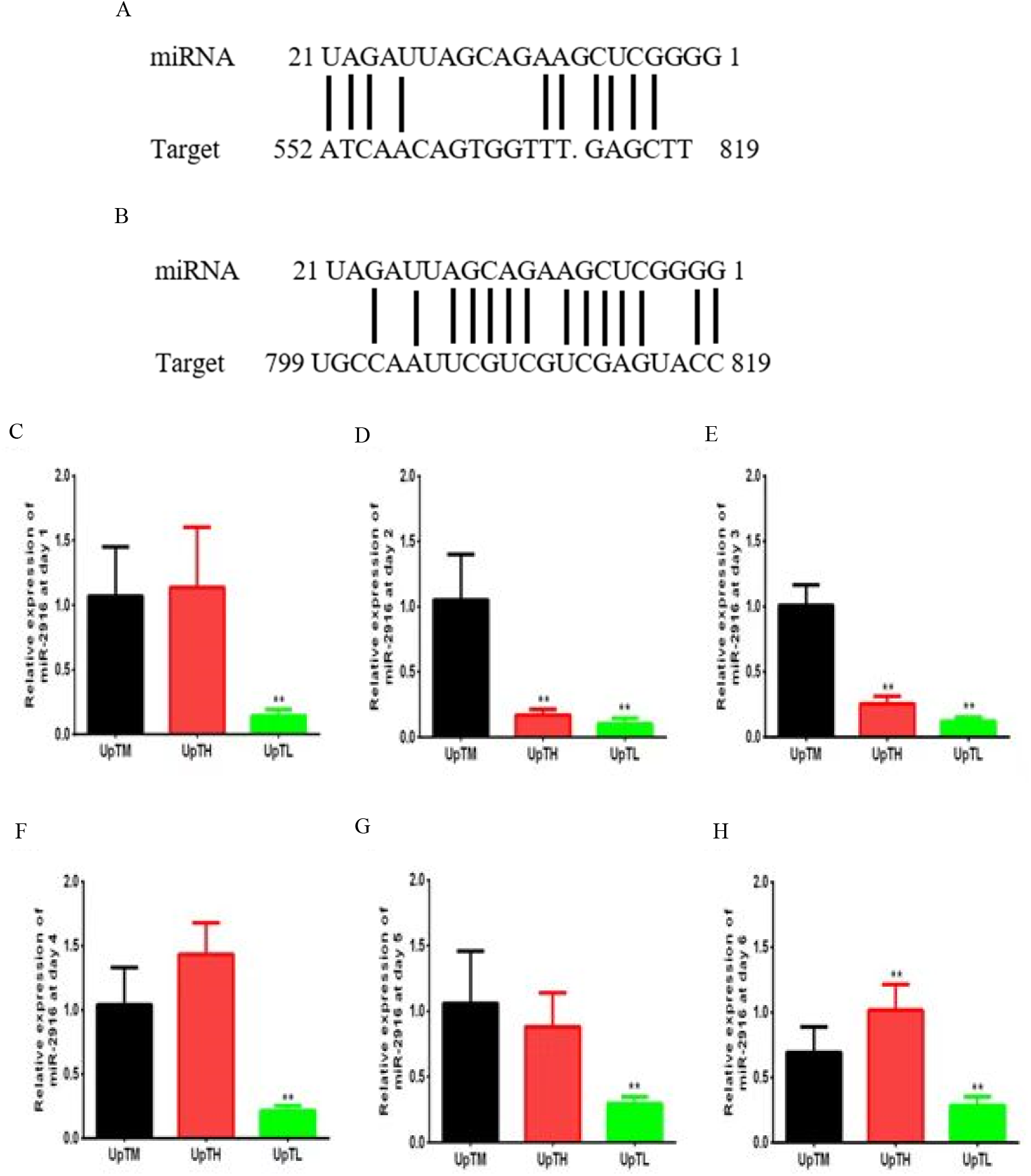
The levels of *miR-2916* in *U. prolifera* cultured under different temperature at different time points. (A) The target position of miR-2916 and PCNA in *U. prolifera*. (B) The target position of miR-2916 of *U. prolifera* and PCNA of *Chlamydomonas reinhardtii*. (C-H) The levels of miR-2916 in *U. prolifera* samples, which were cultured at different temperatures and at different time points. UpTM, protoplasts cultured at 20°C; UpTL, protoplasts cultured at 4°C; UpTH, protoplasts cultured at 36°C; **P<0.01, vs. UpTM groups. The data were analyzed by using one-way ANOVA (Tukey’s test).

### An analysis the interaction between PCNA and CyclinA

To detect the PCNA-binding proteins in the cell lysates of *U. prolifera*, we performed pull-down assays and liquid chromatography/mass spectroscopy (LC-MS/MS). The results shown that there are 58 differential expressed proteins, including 24 up- and 34 down-regulated proteins. Gene ontology (GO) function analysis of 1,069 credible differential proteins showed that 177, 125, and 767 proteins were enriched in MF (Molecular Function), CC (Cell Component), and BP (Biological Process), respectively (Figure 4A). KEGG analysis of 58 credible differential proteins, we found that the ways to enrich the most proteins included that carbon fixation in photo-synthetic organizations, arachidonic acid metabolism, and nicotinate and nicotinamide metabolism (Figure 4B, C). The interaction of different proteins was analyzed by STRING database, and the results showed that the interacting proteins were mainly concentrated in carbon metabolism, citrate cycle, and carbon fixation in photosynthetic organisms (Figure 4D). The results showed that, on the one hand, PCNA participated in the carbon fixation process in photosynthesis by regulating nucleotide metabolism and pentose phosphate cycle pathway; on the other hand, PCNA can also mobilize the activity of enzymes in the antioxidant system, such as phospholipid-peroxide glutathione activity and L-aspartate oxidase activity, to resist temperature stress. Meanwhile, PPI protein interaction network analysis suggested that UpPCNA might bind some heat shock proteins to participate in the molecular process of temperature stress resistance in the *U. prolifera*.

**Figure 4.**
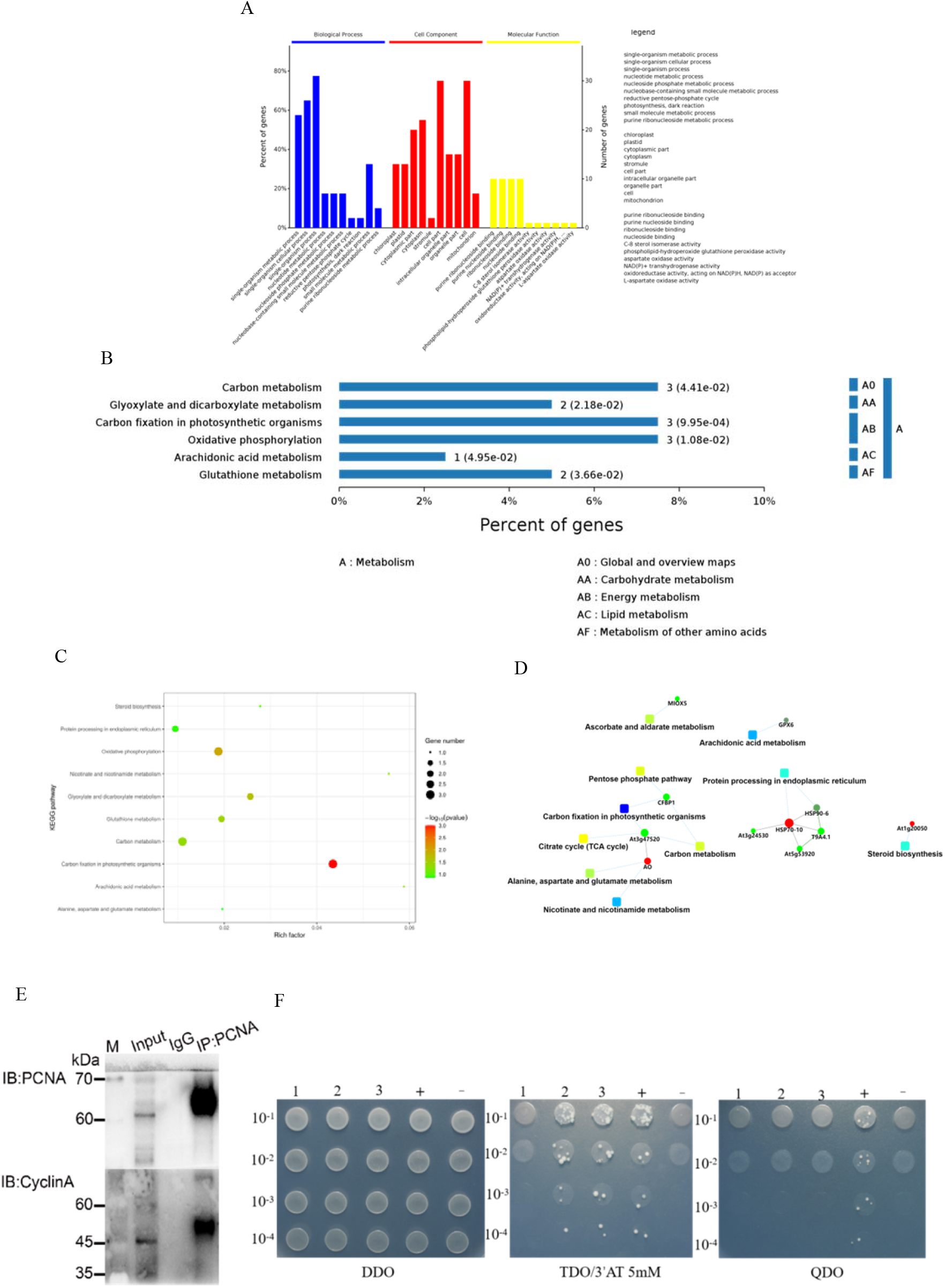
An analysis the interaction between PCNA and CyclinA. (A) GO functional analysis of the differentially expressed proteins. There are 10 significant enrichment categories in BP, CC and MF respectively. P-values is set to 0.05, and the same category is sorted by p value. (B) The annotation for KEGG pathway of differentially expressed proteins. The abscissa represents the proportion of enrichment in each signaling pathway, and the ordinate represents the KEGG term. (C) The bubble picture of KEGG annotation with differentially expressed proteins. The abscissa “Rich factor” represents the enrichment factor, which is defined as the ratio of the protein annotated to the total differentially expressed protein annotated in the pathway. Greater the Rich factor, higher would be the degree of enrichment. The ordinate is the KEGG terms. (D) PPI analysis. (E) Co-Immunoprecipitation assay. IgG, the negative control group; Input, the protein sample without CO-IP treatment. (F) Yeast-two-hybrid assay. 1 : Y2H [PGBKT7-*PCNA*+PGADT7] ; 2: Y2H [PGBKT7-*PCNA*+PGADT7-*CyclinA*];3: Y2H [PGBKT7-*PCNA* + PGADT7-*NPR*]; +: Y2H [Pgbkt7-*53* + Pgadt7-*T*];-: Y2H [Pgbkt7-*lam* + Pgadt7-*T*].

In previous research studies, researchers have detected direct interactions between PCNA and CyclinA. To understand the interacting patterns of PCNA and CyclinA proteins, we investigated the Co-Immunoprecipitation of PCNA and CyclinA. The protein expression of CyclinA and PCNA was detected in the PCNA group and the Input group of PCNA. However, the protein expression of CyclinA was not detected in the IgG group (Figure 4E). The results indicated the binding of CyclinA with PCNA. To determine the potential interaction between PCNA and CyclinA proteins in *U. prolifera*, we next performed a yeast-two-hybrid method. The results showed that only yeast cells were harboring pGBKT7-*PCNA* and pGADT7-*CyclinA*. Hence, they could survive on SD-medium that lacked Leu, Trp, His, and Ade (Figure 4F). Based on these results, we concluded that PCNA interacted with CyclinA in *U. prolifera*.

### High temperature stress inhibited the survival of protoplasts and the formation of cell wall

To determine the survival of protoplasts and the cell wall regeneration capacity in the temperature stresses (UpTH and UpTL), protoplasts were isolated the from *U. prolifera* by enzymatic hydrolysis. As shown in Figure 5A-E, under the action of enzymatic hydrolysate, the cell wall gradually disintegrated, and protoplasts were gradually released. Protoplasts were stained with Evans Blue and Fluorescent Brightener 28 (FB28). The results showed that, because protoplasts had intact cell membranes and had not cell wall components, they could not be stained by Evans Blue and FB28 (Figure 5F-H).

**Figure 5:**
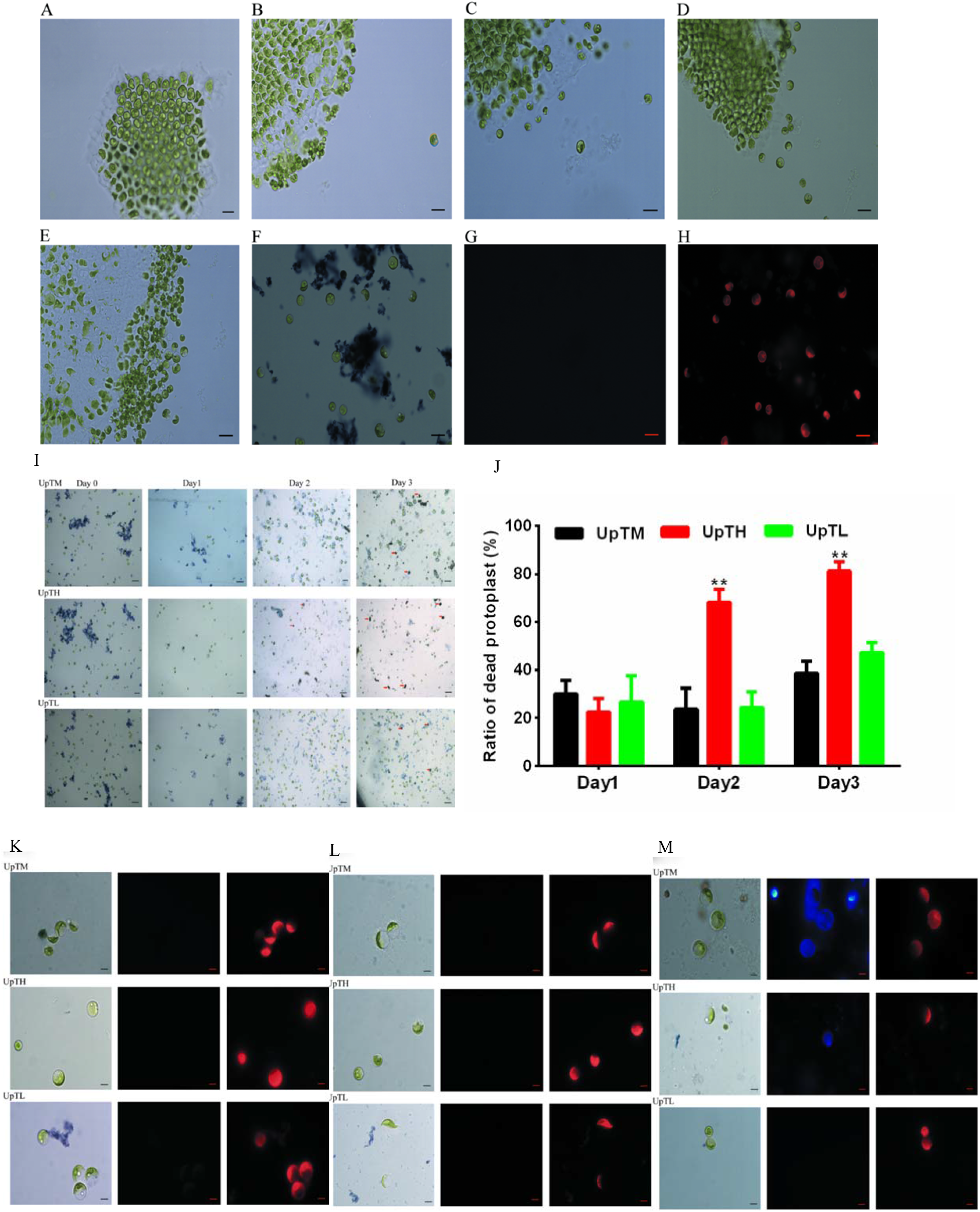
The survival and the formation of cell wall in protoplasts cultured under different temperature at different time points. (A-E) *U. prolifera* was treated with enzyme-digestation buffer for 0 h, 1 h, 2 h, 3 h, 4 h at 25°C. (F) Protoplasts were stained with Evans blue buffer. (G) Protoplasts were stained with Fluorescent Brightener 28 (FB28) buffer. (H) Protoplasts were observed under Lecia fluorescence microscope (EX: 488 nm). Scale bar, 20 μm. (I) The effect of temperature on protoplasts apoptosis was observed by Evans blue staining; Red arrows represent the dead protoplasts, Scale bar 20 μm. (J) The statistical analysis of the rates of dead protoplasts. (K-M) The effect of temperature on protoplast cell wall formation was observed by FB28 staining; A blue protoplast was present within the regenerated cell wall; Scale bars, 5 μm. UpTM, protoplasts cultured at 20°C; UpTL, protoplasts cultured at 4°C; UpTH, protoplasts cultured at 36°C; **P<0.01, vs. UpTM groups. The data were analyzed by performing using one-way ANOVA (Tukey’s test).

Protoplasts in good growth conditions were cultured under temperature stresses (UpTH and UpTL), and stained with Evans Blue or FB28. The results showed the mortality of protoplasts was significantly decreased under the UpTH for 2 days, and the UpTL had not impact on the mortality of protoplasts (Figure 5I). Figure 5K-M showed that the protoplasts were cultured for 3 days at UpTL and UpTM and the synthesis of the cell wall was initiated; however, protoplasts were not stained by FB28 at UpTH. The results indicated that the UpTH inhibited the survival of protoplasts and the formation of cell wall.

### PCNA inhibitor (T2AA) increased the sensitivity of protoplasts under temperature stress

To determine the effect of temperature stresses (UpTL and UpTH) on division of protoplast, protoplasts that entered the cell division and had cell wall were cultured under temperature stresses for 5 days. The results indicated that low temperature stress does not affect the division of protoplast, but high temperature stress stopped the protoplasts’s division (Figure 6A). In addition, the protoplasts were placed at UpTM and treated with different concentrations of T2AA. The culture was maintained for 1 day, 2 days, and 3 days. The results showed that the expression of PCNA was significantly lower in the protoplasts treated with 50 μM T2AA for 2 days compared with control samples (0 μM T2AA) (P<0.01, Figure 6B, C); Meanwhile, the results of Evans blue staining showed that 10 μM T2AA had no effect on the survival of protoplasts, but 50 μM and 100 μM T2AA could cause the death of protoplasts (Figure 6D). This implied that the death of protoplast can be promoted by inhibiting the activity of PCNA.

**Figure 6:**
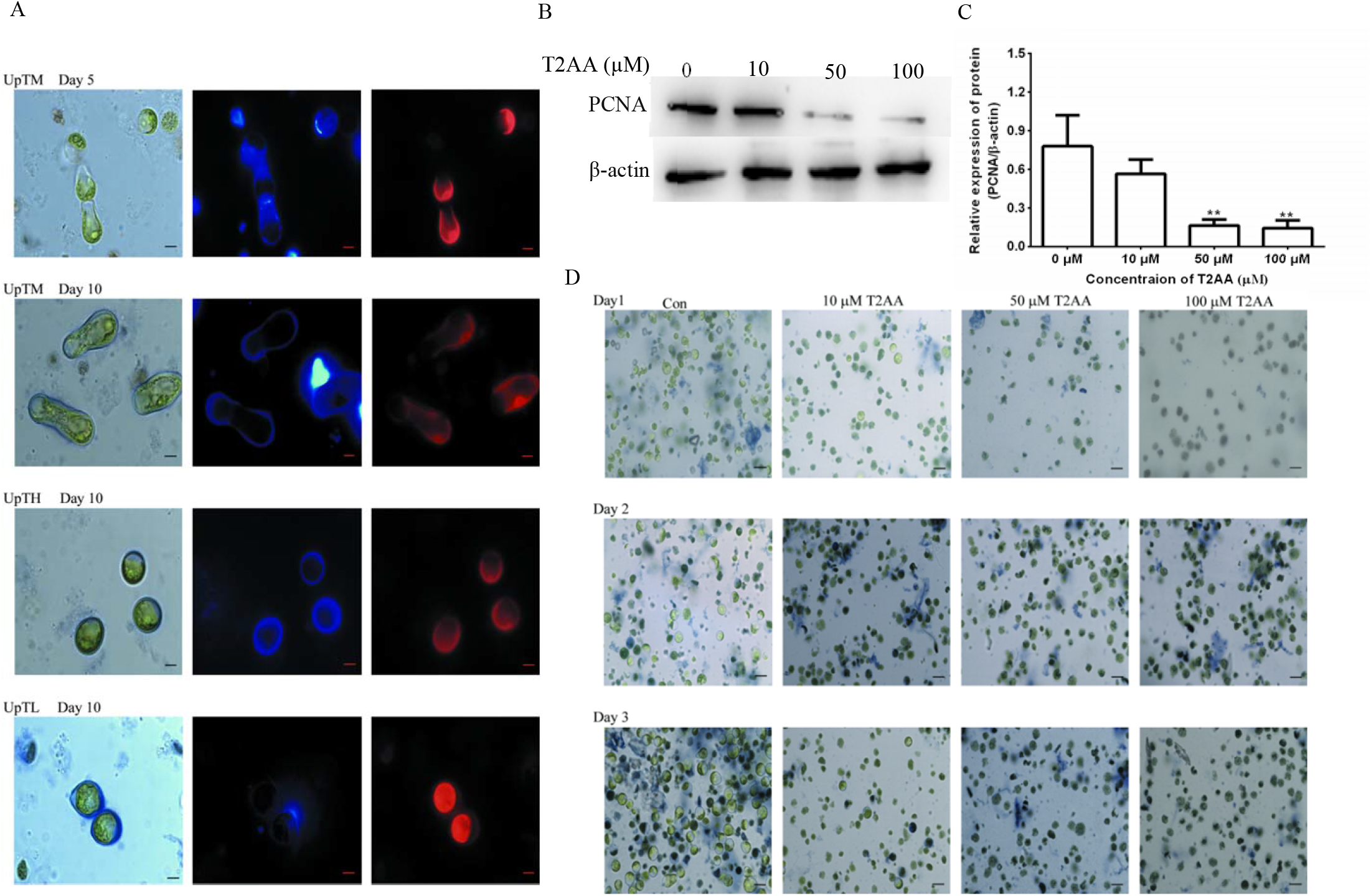
PCNA inhibitor (T2AA) increased the sensitivity of protoplasts under different temperature. (A) The effect of temperature on the division of protoplasts with the cell wall was observed by FB28 staining; Protoplast within the regenerated cell wall is blue; Scale bars, 5 μm. (B-C) The expression of PCNA in protoplasts, which were incubated with different concentrations of T2AA, was verified by performing Western blot analysis. (D) The protoplasts were incubated with different concentrations of T2AA (0 μM, 10 μM, 50 μM, or 100 μM) at 20°C for 1 day, 2 days, and 3 days, respectively. Then, protoplasts were stained with Evans Blue. Scale bars, 20 μm.

## Discussion

*Ulva prolifera*, as a kind of macroalgae, distributed along the coasts of China and many other countries of the world (He et al., 2018a). *U. prolifera* growth is usually affected by different adverse environmental conditions. Among various adverse environmental conditions, temperature is an important factor influencing the growth and reproduction of *U. prolifera*(Li et al., 2012; Wang et al., 2012). In this study, we screened differentially expressed genes (DEGs) related to the cell proliferation of *U. prolifera* under temperature stress, including *PCNA, Cyclins, CDKs*, and *MAPKs*. In plant cells, cyclins (CYC) and cell cycle-dependent kinases (CDKs) jointly control the cell cycle in terms of growth and mitogenic signals. Presently, more than 60 kinds of plant cyclins proteins have been reported, and the most widely investigated are mainly the four families of cyclinA, cyclinB, cyclinC, and cyclinD(Wang et al., 2020). Moreover, PCNA is a highly conserved protein, which acts as a DNA polymerase δ sliding clamp. This protein is present in all the eukaryotes and archaea (Bergink and Jentsch, 2009). Using the transcriptome data, we successfully cloned the full-length cDNAs of *PCNA, CyclinA*, and *UpMAPK17*. By performing BLAST analysis, we confirmed that these genes have a similar expression in plants and algae.

The expression of PCNA, and A-, B-, and D-type cyclins, such as Ntcyc29 (a cyclin B gene) and CycD3-1 was associated with cell proliferation of plants (Zhou et al., 2002). Temperature stresses (UpTL and UpTH) induced the expression of *PCNA* and *CyclinA* of *U. prolifera*. And, CyclinA was identified as a PCNA-interacting protein. As reported, PCNA bound with CyclinA-CDK2 to promote the phosphorylation of the replication factor C (RFC) and DNA polymerases during mammalian cell division and proliferation(Sobczak-Thepot et al., 1993; María de la Paz Sánchez et al., 2002). Hence, PCNA-CyclinA signaling pathway may play an important role in the *U*.*prolifera* regulating the proliferation to acclimatize itself to temperature stress, and may be a potential target for preventing the “green tide”.

Interestingly, temperature stresses (UpTL and UpTH) activated the MAPK signaling pathway. High temperature enhanced the transcription level of *miR-2916* in *U. prolifera*. And, we predicted that miR-2916 has binding sites at positions -552∼-772 of PCNA. The MAPKs and microRNAs (miRNAs) have been shown to play a central role in the control of cell proliferation and temperature tolerance(Voinnet, 2009; Hackenberg et al., 2015; Niu et al., 2016; Banerjee et al., 2020). As is well known, PCNA can bind to CDK2-CyclinA complex; Then, the MAPK signaling pathway is activated by the phosphorylated CDK2 complex to regulate cell cycle and temperature tolerance(Sasabe and Machida, 2012). Previous studies have found that miR-843 participated in the resistance process of *U. prolifera* to nitrogen stress by regulating the expression of Cyclins (Yang et al., 2013). Therefore, temperature stress may affect cell proliferation of *U. prolifera* through miR-2916-PCNA-CyclinA-MAPK mediated pathway: temperature stress induced the expression of *PCNA* and *miR-2916*, and promoted the binding of PCNA to CyclinA; The combination of PCNA and CyclinA relieved the inhibitory effect of CyclinA complex on MAPK signaling pathway. The activated MAPK signaling pathway promoted the expression of a series of downstream signaling molecules which were related to cell proliferation and division. However, what are the downstream genes of MAPK signaling pathway in *U. prolifera*, and which molecules are phosphorylated by CyclinA to activate MAPK signaling pathway? These still need to be further discussed in later research.

In addition, our study suggested that low temperature had no obvious effect on the survival, the formation of cell wall, and the division of protoplasts; However, high temperature inhibited the survival of protoplasts, the formation of cell wall of protoplasts, and the division of protoplasts. The results indicated that the molecular mechanism of *U. prolifera* in oversummering and overwintering may be different. The results consisted with the speculations of algae experts on the source of *U. prolifera* (Liu et al., 2012). The results confirmed the view that *U. prolifera* might come from microscopic propagules. The microbreeders couldn’t survive in the high temperature environment, but can divide and proliferate under the low temperature conditions. The proliferation rate is slow under the low temperature, but the appropriate temperature in April to June can provide hotbed for the asexual reproduction and sexual reproduction of *U. prolifera*. PCNA inhibitor decreased the protoplast activity of *U*.*prolifera*. Thus, once the expression of *PCNA* was inhibited, protoplast could not continue cell division, which provided a certain solution strategy for the control of *U. prolifera*.

## Materials and Methods

### Alga material and treatment

*Ulva prolifera* materials were collected from the coast of Qingdao (36°48′39.75′’N; 121°38′10.88′’E), China. *U. prolifera* were cultured in fresh distilled seawater supplemented with macro-elemental solution as previously reported (He et al., 2018b) and at 20°C in SPX-GB-250 intelligent illumination incubators (Botai, Shanghai, China). Light was provided by a halogen lamp at a parabolic aluminized reflector of 100 μmol/(m·s) for 12 h. The cultured seawater was renewed every 2 d, and temperature stress experiments were performed after pre-treatment. Cool-white fluorescent light was provided on a 12 L:12 D cycle. There were three temperature regimes: 4°C was set as the UpTL group, 36°C was set as the UpTH group, and 20°C was set as the UpTM group.

### Library preparation and sequencing

The cDNA library was constructed using TruSeqTM RNA sample preparation Kit from Illumina (San Diego, CA). After quantified by TBS380, the resulting cDNA library was then sequenced using the Illumina HiSeq Xten/NovaSeq 6,000 sequencer (2 × 150 bp read length) by Shanghai MajorBio Co., Ltd.

### Differential expression analysis and Functional enrichment

To determine the expression level of each transcript, we used the method of fragments per kilobase of exon per million mapped reads (FRKM). The abundance of genes was quantified by performing the technique of RNA-Seq by Expectation-Maximization (RSEM) (http://deweylab.biostat.wisc.edu/rsem/)(Li and Dewey, 2011). To perform differential expression analysis, we used the statistical software of R EdgeR (Empirical analysis of Digital Gene Expression in R, (http://www.bioconductor.org/packages/2.12/bioc/html/edgeR.html)(Robinson et al., 2010). ene ontology (GO) database, and the Kyoto Encyclopedia for Genes and Genomes (KEGG) were used to perform functional enrichment analysis. In GO terms, we identified the significantly enriched DEGs. Unlike the background of whole-transcriptome, some metabolic pathways were associated with the Bonferroni-corrected P-value ≤0.05. In this experiment, GO functional enrichment and KEGG pathway analysis were carried out by using Goatools (https://github.com/tanghaibao/Goatools) and KOBAS (http://kobas.cbi.pku.edu.cn/home.do)(Xie et al., 2011).

### Co-immunoprecipitation (Co-IP) assay

PCNA antibody (1:100) was coupled to Protein A+G agarose beads. Then, PCNA antibody-resin mixtures were incubated with supernatants of cell lysates. Finally, elution buffer eluted the Co-immunoprecipitated proteins. The eluates and the samples were stored at -80°C temperature for further analysis.

### LC-MS/MS

The sample was digested, desalted, and dissolved in the sample solution containing 0.1% formic acid and 2% acetonitrile. Then, the mixture was vortexed thoroughly and centrifuged. The supernatant was loaded onto a 150 μm×15 cm in-house column Ultimate 3,000 series of HPLC system was coupled to a Q Exactive™ Hybrid Quadrupole-Orbitrap™ Mass Spectrometer via a nano-electrospray source. The separated eluents were discharged from the column by using a 60 min gradient of 6– 95% of buffer B (80% ACN, 0.1% formic acid), which was added at a flow rate of 600 nL/min. Each sample (5 μL) was separated in a binary phase, which consisted of the mobile phase A (0.1% formic acid in water) and mobile phase B (0.1% formic acid in water-80% acetonitrile).

The raw MS files were analyzed, and a protein database was searched to identify the species of the samples. MaxQuant softwareversion1.6.2.10 was used for analysis. The main parameters were as follows: protein modifications: carbamidomethylation (C) (fixed), oxidation (M) (variable), and Acetyl (Protein N-term) (variable); the specificity of the enzyme to trypsin; the permission of two missed cleavages; the tolerance of the precursor ion mass was 20 ppm, and the MS/MS tolerance was 20 ppm. We selected only the peptides identified with high confidence. Then, we performed downstream protein identification analysis on these peptides.

### Protoplast isolation, fluorescent brightener 28 (FB28) and Evans blue staining

Protoplasts were prepared with the methods already reported(Yang et al., 2021). Thereafter, protoplasts were placed in light incubators, which were maintained at 4°C, 20°C, or 36°C for 1 day, 2 days, or 3 days respectively. Protoplasts were stained with 1% Evans Blue dye to confirm the viability of them. Then, we observed the protoplasts with a UV florescence microscope (Leica, DM2500, Wetzlar, Germany). At the same time, the protoplasts were stained with 0.1% (w/v) FB28 solution (SigmaAldrich, St. Louis, MO, USA) to determine the regeneration capacity of protoplasts.

## Acknowledgements

This work was supported by the National Key R & D Program of China (No. 2016YFC1402102), National Nature Science Foundation of China (No. 41976109), the MNR Key Laboratory of Eco-Environmental Science and Technology of China (MEEST-2020-2), Chinese Postdoctoral Science Foundation (No. 2020M681698), Natural Science Foundation of Jiangsu Province (BK20200882), Jiangsu Planned Projects for Postdoctoral Research Funds (No. 2020Z300), Natural Science Foundation of the Jiangsu Higher Education Institutions of China (No. 20KJD170004), National Science Foundation Award (No. 42106099), and the Project Funded by the Priority Academic Program Development of Jiangsu Higher Education Institutions (PAPD).

## Author contributions

H.Y.H., J.J.Y., and Y.H. designed the study, performed major experiments, and wrote the manuscript. Z.Y.L., C.W.F., and D.R.Z performed investigation and analyses. M.R.L., A.M.L., and J.W.D. performed specific experiments. J.S.L. and H.Y.G. performed data curation. S.D.S. performed project administration and supervised. All authors read and approved this paper. H.Y.H., J.J.Y., and Y.H. contributed equally to this work.

## Data availability

The data that support the findings of this study are openly available from the corresponding author upon reasonable request.

